# Bayesian inference of transcription dynamics from population snapshots of single-molecule RNA FISH in single cells

**DOI:** 10.1101/109603

**Authors:** Mariana Gómez-Schiavon, Liang-Fu Chen, Anne E. West, Nicolas E. Buchler

## Abstract

Single-molecule RNA fluorescence *in situ* hybridization (smFISH) provides unparalleled resolution on the abundance and localization of nascent and mature transcripts in single cells. Gene expression dynamics are typically inferred by measuring mRNA abundance in small numbers of fixed cells sampled from a population at multiple time-points after induction. The sparse data that arise from the small number of cells obtained using smFISH present a challenge for inferring transcription dynamics. Here, we developed a computational pipeline (BayFish) to infer kinetic parameters of gene expression from smFISH data at multiple time points after induction. Given an underlying model of gene expression, BayFish uses a Monte Carlo method to estimate the Bayesian posterior probability of the model parameters and quantify the parameter uncertainty given the observed smFISH data. We tested BayFish on smFISH measurements of the neuronal activity inducible gene *Npas4* in primary neurons. We showed that a 2-state promoter model can recapitulate *Npas4* dynamics after induction and we inferred that the transition rate from the promoter OFF state to the ON state is increased by the stimulus.

**Author Summary:** Gene expression can exhibit cell-to-cell variability due to the stochastic nature of biochemical reactions. Single cell assays (e.g. smFISH) directly quantify stochastic gene expression by measuring the number of active promoters and transcripts per cell in a population of cells. The data are distributions and their shape and time-evolution contain critical information on the underlying process of gene expression. Recent work has combined models of stochastic gene expression with maximum likelihood methods to infer kinetic parameters from smFISH distributions. However, these approaches do not provide a probability distribution or likelihood of model parameters inferred from the smFISH data. This information is useful because it indicates which parameters are loosely constrained by the data and suggests follow up experiments. We developed a suite of MATLAB programs (BayFish) that estimate the Bayesian posterior probability of model parameters from smFISH data. The user specifies an underlying model of stochastic gene expression with unknown parameters (*θ*) and provides smFISH data (*Y*). BayFish uses a Monte Carlo algorithm to estimate the Bayesian posterior probability *P*(*θ|Y*) of model parameters. BayFish is easily modified and can be applied to other models of stochastic gene expression and smFISH data sets.

## Introduction

Cell-to-cell variation in gene expression across an isogenic population is a fact of life [1]. The initiation of transcription involves a series of stochastic biochemical events, including the binding of transcription factors and RNA polymerase to the promoter of a gene [2]. Distinct promoter states often arise when one of these biochemical events is rate-limiting. The existence of multiple promoter states with different expression rates can generate transcriptional bursting, which are episodes of transcriptional activity followed by long periods of inactivity [3]. This phenomenon has been observed in bacteria [4], yeast [5, 6], fly [7] and mammals [8].

Cell-to-cell variability in gene expression is studied using experimental techniques that measure transcription levels in single cells [4, 9–14]. One such technique, single-molecule RNA fluorescence *in situ* hybridization (smFISH), measures the abundance and localization of individual transcripts in single cells. We have used smFISH to measure transcripts of the neuronal activity inducible gene *Npas4* in primary neurons after membrane depolarization with elevated extracellular potassium (Fig. 1). Each individual transcript is bound by fluorescent DNA probes and appears as a bright, diffraction-limited spot in a fluorescence microscope [9, 15]. In cases where there are multiple transcripts (e.g. active transcriptional sites at gene loci), the measured intensity is significantly brighter. Our *Npas4* smFISH measurements showed a surprising amount of cell-to-cell variation in both transcript levels and active gene loci given that all neurons were exposed to a uniform external stimulus. Given prior studies of cell-to-cell variability in gene expression in other systems, we infer that this variability in the transcriptional response of activity-inducible genes is likely to arise from the probabilistic activation of transcriptional bursting at single alleles. We thus reasoned that we could use our single cell transcriptional variability to build a model of activity-inducible *Npas4* induction that would inform our quantitative understanding of the transcriptional processes that drive dynamic changes in *Npas4* expression following neuronal activation.

**Fig 1.**
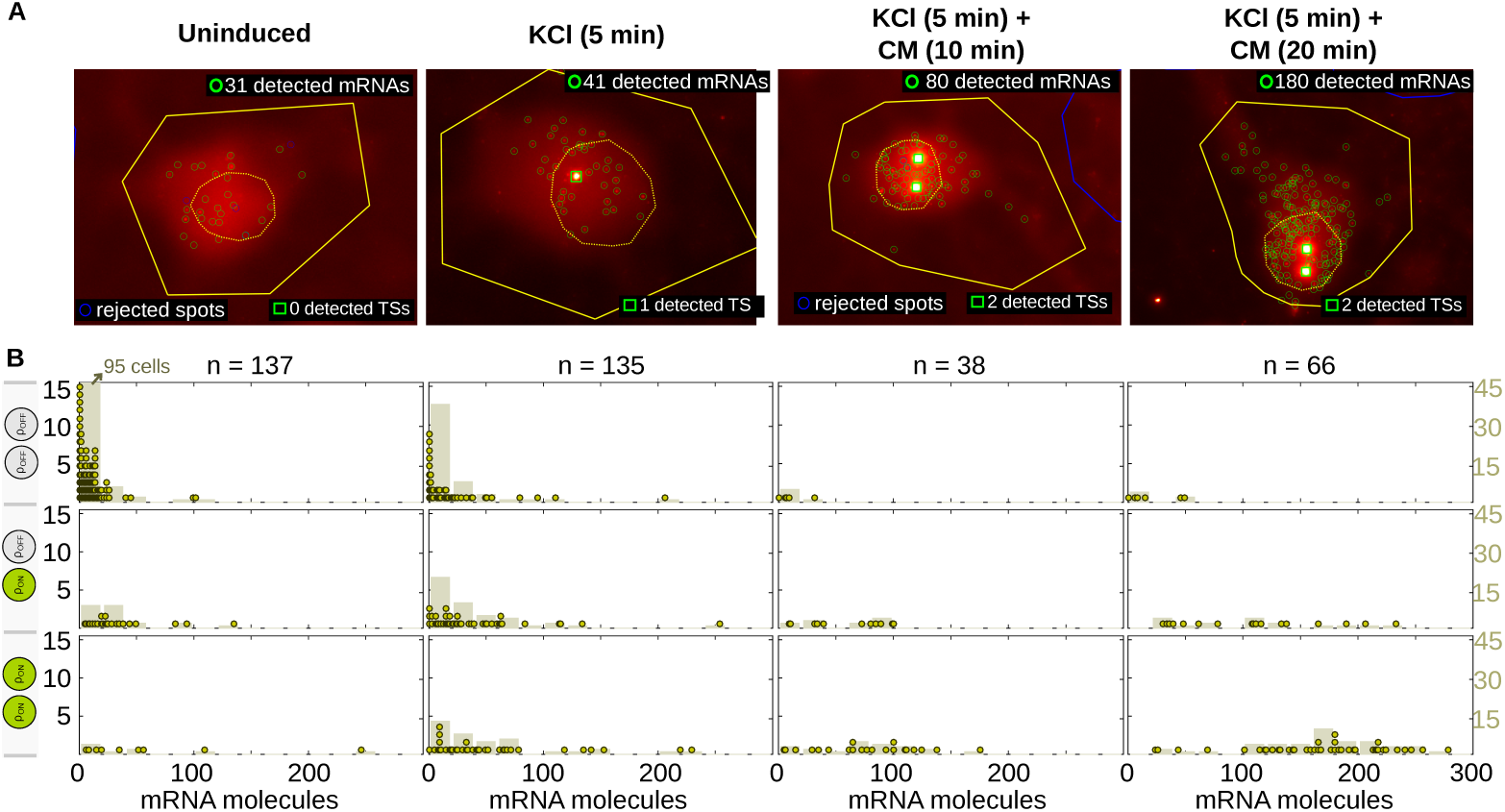
Single-molecule fluorescennce *in situ* hybridization data of *Npas4* mRNA in primary neurons after membrane depolarization. Measurements are shown before the stimulus (uninduced), 5 minutes after KCl exposure, and an additional 10 and 20 minutes later after cells were returned to conditioned medium (CM); see *Methods*. (A) Example of an image-processed cell at each time point. We show detected mRNAs (green circles) and active transcription sites (TSs; green squares) within the cell contour (yellow line) and nucleus (dashed yellow line). Neurons are post-mitotic and, thus, we observed up to two active gene loci per diploid cell. (B) For each condition, the histogram of the number of mRNA molecules binned by the number of TSs (dots, left y-axis). Smoothed histogram with bins of 20 mRNAs (bars, right y-axis). The total number of cells (*n*) per time sample is listed at the top.

To better understand the origins of transcriptional bursting in this immediate-early gene, we combined a mathematical model of stochastic gene expression and our smFISH data to infer which model parameters are regulated by the stimulus. The challenge is that we had ~ 100 cells per time point (Fig. 1). The low number of measured cells means that the observed frequency distribution of transcripts is sparse (i.e. many zero entries) and inferred model parameters will be sensitive to sampling error. This is a common problem for studies using primary cells where it is challenging to routinely generate and analyze massive amounts of smFISH data. To address this challenge, we developed a computational pipeline (BayFish) that infers the best model parameters from sparse smFISH data and rigorously quantifies the uncertainty in those parameters. We used BayFish on our *Npas4* smFISH data to infer the parameters of an underlying 2-state model of gene expression that were likely affected by the stimulus.

## Method

BayFish is a Monte Carlo method that estimates the Bayesian posterior probability *P*(*θ|Y*) of model parameters (*θ*) given the observed smFISH data (*Y*) at different time points before and after induction. Bayes theorem states *P*(*θ|Y*) = *P*(*Y θ*) *P*(*θ*)*/P* (*Y*) where *P*(*Y θ*) is the likelihood 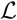 of the data given the parameters. *P*(*θ*) and *P*(*Y*) are the prior probability distributions of parameters and data, respectively. Each iteration of the Monte Carlo method uses several numerical sub-routines to (1) calculate the time evolution of the mRNA distribution given a set of model parameters (*θ*), (2) evaluate the likelihood that the smFISH data (*Y*) were sampled from this distribution, or 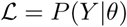, and (3) calculate the Bayesian posterior probability 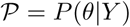 given the likelihood and priors. The global program is based on the Metropolis Random Walk algorithm [16, 17]:

1. Specify a mathematical model of stochastic gene expression that has an unknown set of parameters *θ*.
2. Choose an initial *θ* and calculate the corresponding likelihood 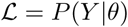 and Bayesian posterior probability 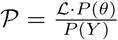 using several numerical sub-routines.
3. Iterate over *t* = {1, 2, …, *T*} as follows:

a. Draw a random proposal 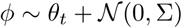, where 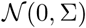 is a Multivariate Normal distribution with the same dimension as *θ*, zero mean and Σ covariance matrix.
b. Evaluate the likelihood of the proposal 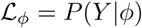 using several numerical sub-routines.
c. Calculate the Bayesian posterior probability 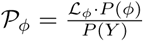.
d. Update parameters *θ*_*t*+1_ *← φ* and 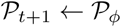 with probability min 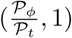; otherwise, *θ*_*t*+1_ *← θ*_*t*_ and 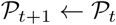.

Over time, the algorithm will generate a Markov chain of *θ*_*t*_ whose distribution converges to the Bayesian posterior probability *P*(*θ|Y*). BayFish saves the likelihood 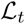 and *θ*_*t*_ of each step. After discarding the early part of the chain (the “burn-in” phase), the remaining *θ*_*t*_ were used to estimate the Bayesian posterior probability *P*(*θ|Y*); see *Methods*. Below, we explain and justify the sub-routines of our pipeline.

### Mathematical model of stochastic gene expression

We considered a 2-state model of gene expression (Fig. 2), where each promoter can be in an inactive OFF state with a basal transcription level (synthesis rate *µ*_0_) or an active ON state with a higher transcription level (synthesis rate *µ*_1_). Transitions between promoter states occur with a promoter activation rate *k*_1_ and a promoter deactivation rate *k*_0_. We chose a 2-state model because it is the simplest model that can generate transcriptional bursting, a feature observed in our smFISH data. Each promoter allele was assumed to be regulated independently of the other [18], but other scenarios could be implemented as needed. The 2-state model parameter set, which determines the dynamics of mRNA and active promoters, is *θ* = *µ*_0_, *µ*_1_, *k*_1_, *k*_0_.

**Fig 2.**
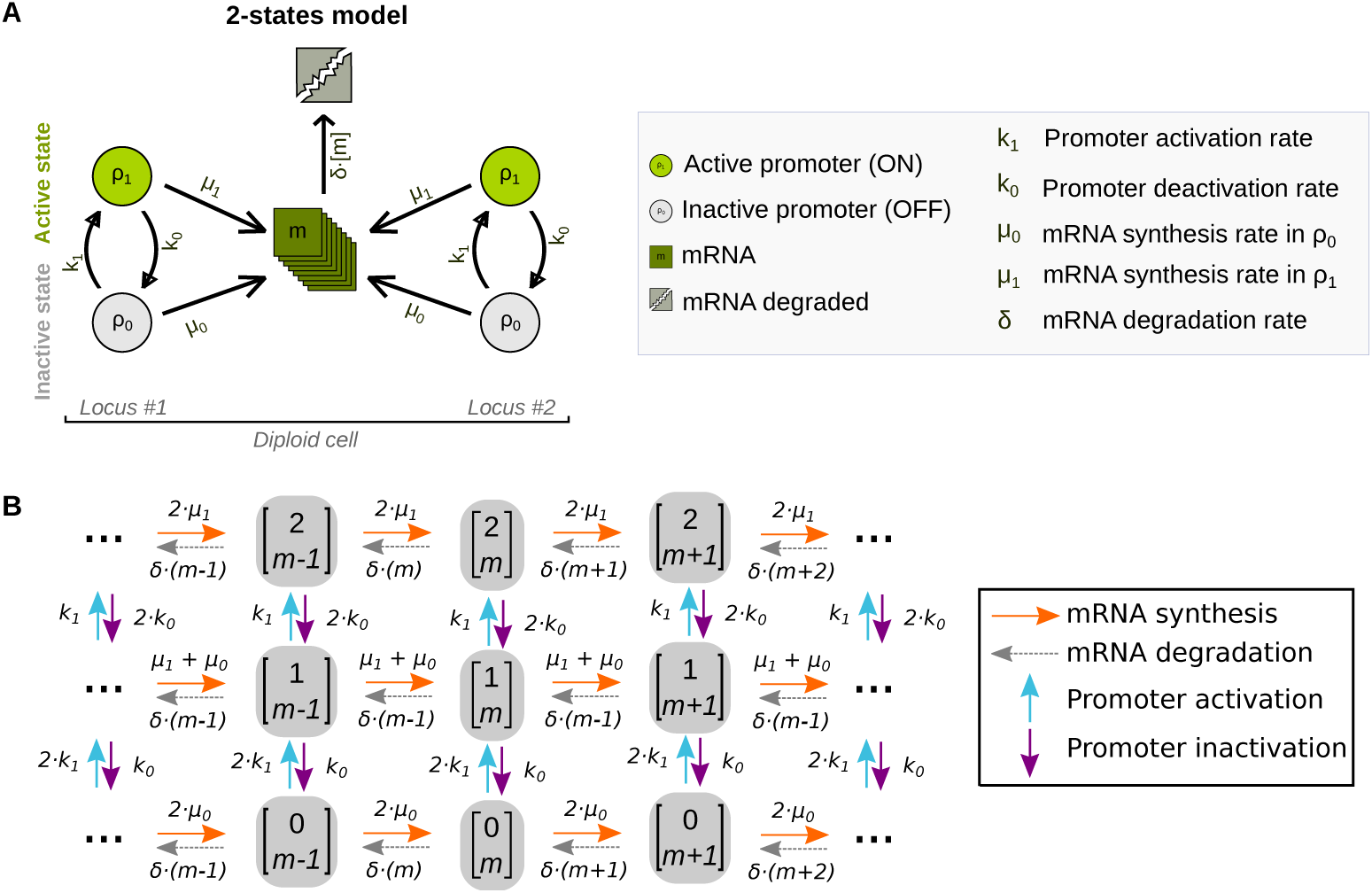
A 2-state model of gene expression. (A) Each diploid cell has two genetic loci and the promoter (circle) of each gene can be either in an active (*ρ*_1_) or inactive (*ρ*_0_) state. Each gene synthesizes mRNA molecules (*m*) with rate *µ*_1_ or *µ*_0_ if the promoter is active or inactive, respectively. Transitions between promoter states occur with a promoter activation rate *k*_1_ and a promoter deactivation rate *k*_0_. Each mRNA is degraded with rate *δ*. The mRNA degradation rate constant of *Npas4* has been measured [19] and was fixed to *δ* = 0.0559 min^*−*1^. (B) Possible biochemical reactions and cell states of our model. A cell state **x** (grey box) is the number of active promoters *ρ*_1_ ∈ 0, 1, 2 and mRNA molecules *m* ∈ 0, 1, 2*, . . . , M* in a cell, or **x** = [*ρ*_1_, *m*]^*T*^. There are four possible biochemical reactions that change a cell from one state to another state: (1) Promoter activation (blue arrow), which increases *ρ*_1_ by one; (2) promoter inactivation (purple arrow), which decreases *ρ*_1_ by one; (3) mRNA synthesis (orange arrow), which increases *m* by one; and (4) mRNA degradation (gray arrow), which decreases *m* by one. The propensity or probability per unit time (*a*_*k*_) for a particular reaction (*k*) to occur is listed above the reaction arrows. The propensities depend on the model parameters *θ* = {*µ*_0_, *µ*_1_, *k*_1_, *k*_0_}.

Our smFISH experiments measured gene expression both before and after stimulus. We presumed that gene expression before stimulus was at a steady state determined by one set of model parameters (*θ*_*U*_, unstimulated parameter set). Upon induction, the stimulus changed one or more of the model parameters (*θ*_*S*_, stimulated parameter set). Thus, the distribution of mRNA and active promoter states will evolve towards a new steady-state in response to the changed parameters. Below, we describe how we calculated the stationary distribution of mRNA and active promoters before stimulus using *θ*_*U*_ and how we then calculated the time-evolution of the distribution after stimulus using *θ*_*S*_.

### Time-evolution of the probability distribution

The Chemical Master Equation (CME) is an infinite set of coupled differential equations that describe the dynamics of the probability of the biochemical system being in a particular state ***x*** at time *t*, *P*(***x**, t*) [20, 21]. The probability flow into and out of each state ***x*** is given by:

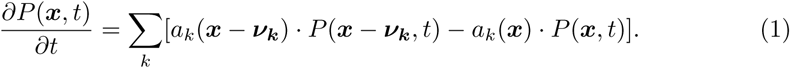

The summation is over all possible biochemical reactions *k* into and out of state **x**:

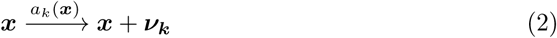

where *a*_*k*_(***x***) *∂t* is the probability that the biochemical reaction *k* will occur within the infinitesimal time interval *∂t* given that the system is in state ***x***. The model parameters *θ* affect the propensities of different biochemical reactions (Fig. 2), and the stoichiometric vector 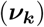 of reaction *k* describes how the system state changes when the reaction *k* occurs. More generally, the CME is written in matrix form:

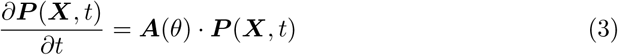

where all possible cell states ***X*** are enumerated as a vector [***x***_**1**_, ***x***_**2**_, . . . , ***x***_***N***_]^*T*^, ***P*** (***X**, t*) is the probability density state vector [*P*(***x***_**1**_, *t*), *P*(***x***_2_, *t*), . . . , *P*(***x***_***N***_, *t*)]^*T*^ of possible states organized identically to ***X***. The state reaction matrix ***A***(*θ*) has elements:

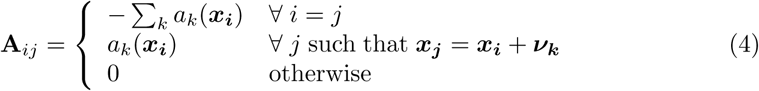

### Pre-stimulus stationary distribution

We assumed that the pre-stimulus distribution of mRNAs and active promoters **P**^*∗*^(***X***) is time-independent and stationary. We calculated the stationary distribution by setting Eq. 3 to zero and determined the nonzero eigenvector **V ≥ 0** in the kernel of **A**(*θ*_*U*_) using the Arnoldi iteration algorithm [22] (*eigs* MATLAB function). Each element of **P**^*∗*^ is given by:

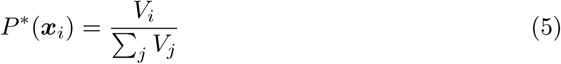

where *V*_*i*_ is the *i*th element in the vector ***V*** = [*V*_1_, *V*_2_, . . . , *V*_*N*_]^*T*^ and Σ_*i*_ *P*^∗^(***x***_***i***_) = 1. The size (*N*) of the vector and matrix is determined by *N* = 3(*M* + 1), where *M* can be infinite. For practical purposes, we chose *M* = 500 because it is finite and larger than the expected mRNA levels in our smFISH data.

### Post-stimulus distribution dynamics

Given an initial distribution ***P*** ^∗^(***X***) at time zero and post-stimulus state reaction matrix ***A***(*θ*_*S*_), the post-stimulus distribution ***P*** (***X**, τ*) at time *τ* after stimulus is:

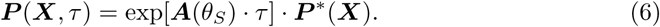

We calculated ***P*** (***X**, τ*) after induction using the same MATLAB routines from the Finite State Projection (FSP) method [23]. We used FSP to verify that our estimated probability distributions for finite *M* were below error threshold (*ϵ* ≤ 10^−12^).

### Likelihood of smFISH data from probability distributions

The smFISH data is a sample of cells at several time points {0*, τ*_1_, *τ*_2_, . . . *τ*_*S*_} after induction. Each cell was in a state contained within [***x***_1_, ***x***_2_, . . . , ***x***_***N***_]^*T*^. The smFISH data vector ***Y*** ^***t***^ for sample *t* is a count of observed cell states, where [*n*_1_, *n*_2_, . . . *n*_*N*_]^*T*^. The likelihood of having sampled the observed data given the calculated distributions ***P*** (***X**, τ*) for model parameters *θ* is a product of multinomial distributions:

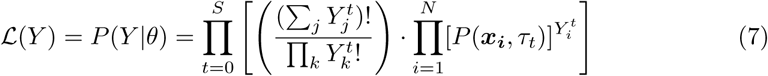

### Calculate the Bayesian posterior probability

The Bayesian posterior probability is the likelihood 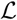 multiplied by *P*(*θ*) and divided by *P*(*Y*), which are the prior probability distributions of parameters and data. These priors are often unknown and *P*(*θ*) and *P*(*Y*) are presumed flat and constant, i.e. any parameter set and data set is equally likely. BayFish assumes flat priors unless specified otherwise. We implemented a Heaviside step function for *P*(*θ*), where the prior was zero for non-physiological parameters (i.e. negative numbers, where any parameter is below 10^*−*8^), but otherwise flat and constant.

## Results

We considered several models where the stimulus can affect multiple parameters. We start by showing the best-fitting (*k*_1_, *k*_0_)-stimulus model where the promoter activation and deactivation rates respond to stimulus, i.e. all other parameters are identical between the pre- and post-stimulus conditions. We ran three replicas of BayFish with different initial parameters for *T* = 10^5^ iterations. If the *Npas4* smFISH data were too few to constrain the model, then we expect the Bayesian posterior distributions to be flat. However, all BayFish replicas converged to identical, well-defined Bayesian posterior distributions of model parameters, which demonstrates that our sparse smFISH data do constrain the parameters of the underlying model (Fig. 3).

**Fig 3.**
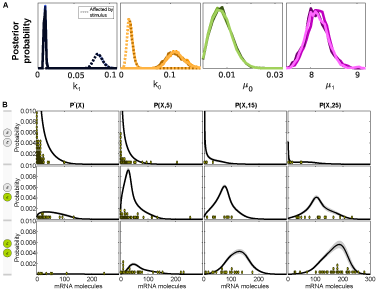
Bayesian posterior distribution of parameters for (*k*_1_, *k*_0_)-stimulus model. (A) Marginal posterior distributions of parameters for BayFish replicas. There are two distributions for *k*_0_ and *k*_1_, one of which is the pre-stimulus parameter (continuous lines) and the other is the post-stimulus parameter (dotted lines). (B) The mean mRNA and active promoter distributions *(**P*** 〈***X**, τ*)〉 as inferred from the Bayesian posterior distribution of parameters. The standard deviation (*σ*_**P**_) is shown in gray. The histogram of experimental data is shown for comparison (green dots).

### Comparing different stimulus models

We systematically considered other parameter combinations that could be affected by the stimulus: *k*_1_-, *k*_0_-, *µ*_1_-, (*k*_1_, *µ*_1_)-, (*k*_0_, *µ*_1_)-, and (*k*_1_, *k*_0_, *µ*_1_)-stimulus models. A one parameter-stimulus model has 5 free parameters and a three parameter-stimulus model has 7 free parameters. It is well-known that models with more parameters have a higher likelihood of fitting the data. To this end, we used several likelihood-based metrics to evaluate different models and penalize those with increasing free parameters (see Methods). These metrics are the *Bayesian Information Criterion* (BIC) [24] and the *Akaike Information Criterion* (AIC) [25], which are based on the maximum likelihood calculated by BayFish. The *Deviance Information Criterion* (DIC) [26] uses both the likelihood and the Bayesian posterior distribution calculated by BayFish.

We ran three replicas of BayFish with different initial parameters for each stimulus model. The three metrics gave identical results to one another (Fig. 4). The Bayesian posterior distributions of parameters for each stimulus model are shown in S1 Fig. The best model with the fewest parameters was the (*k*_1_, *k*_0_)-stimulus model. However, not all parameters are equivalent. Fig. 4 demonstrates that regulation of *k*_1_ by the stimulus consistently gives a better fit to the observed data than regulation by *k*_0_ or *µ*_1_ alone or in combination.

**Fig 4.**
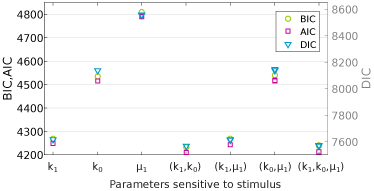
Comparing different stimulus models. We applied Bayesian Information Criterion (BIC), Akaike Information Criterion (AIC), and Deviance Information Criterion (DIC) metrics to the BayFish results obtained from different parameter-stimulus models listed in the *x*-axis. Models with lowest BIC and AIC scores (left, y-axis) and DIC (right, y-axis) are considered to be the most informative models with the fewest parameters. For each stimulus model, three replicas of BayFish were run with different initial conditions; the maximum likelihood observed in the three replicas was used for BIC and AIC metrics, and the full likelihood and Bayesian posterior distribution excluding the “burn-in” period were used for DIC.

## Discussion

Single cell measurements of transcript abundance using smFISH have been combined with mathematical models of stochastic gene expression to elucidate mechanisms of transcriptional bursting [6, 27]. Distributions of mRNAs derived from 2-state promoter models were fit to smFISH data to infer kinetic parameters. These early models presumed that gene expression was at steady-state. More recent papers have used the Finite State Projection algorithm to calculate the time-evolution of promoter-state and mRNA distributions of more complex models (e.g. 3-state promoters) not necessarily at steady state (e.g. after induction) [18, 28]. There is no software package where one can specify a complex model of stochastic gene expression, evaluate the time-evolution of promoter-state and mRNA distributions after induction, and robustly infer parameters from measured smFISH data using the Bayesian posterior distribution.

We developed a suite of MATLAB programs (BayFish) that use Bayesian inference to robustly estimate model parameters from smFISH data. The user specifies a mathematical model of stochastic gene expression with an unknown set of parameters (*θ*) and provides smFISH data (*Y*) at different time points before and after induction. BayFish uses a Monte Carlo method to estimate the Bayesian posterior probability *P*(*θ|Y*) of the model parameters, which elucidates the best-fitting parameters and quantifies their uncertainty. BayFish can be modified to include more complex models of gene expression and different data sets. Bayesian inference is especially useful for experimental systems with smaller smFISH data sets that have large sampling error. As a test case, we used BayFish to extract meaningful biological information from *Npas4* gene expression in single neurons (Fig. 1). We ran BayFish with different variations of the 2-state model and used different Information criteria to infer that the stimulus likely regulates the promoter activation rate (*k*_1_). Future experiments will address mechanisms of activation and cell-to-cell variability in *Npas4* and other immediate-early genes of primary neurons. This can be done by combining genetic and pharmacological perturbations of gene expression with downstream BayFish analysis of multi-color smFISH distributions of several immediate-early genes.

## Methods

### *Npas4* smFISH measurements in single neurons

Neuron-enriched cultures were generated from the cortex of male and female E16.5 CD1 mouse embryos (Charles River Laboratories Inc., Wilmington, MA, USA) and cultured as previously described [29]. Neurons were treated with 1 *µ*M sodium channel inhibitor TTX (Tocris Cookson, Ballwin, MO, USA) at DIV6 and depolarized by elevating extracellular potassium concentration to 55 mM with a isotonic KCl solution at DIV7 [30], which activates L-type voltage-gated calcium channel dependent transcription of *Npas4* [31]. Cells were fixed at 4 time points: no KCl, 5 mins KCl treatment, 5 mins KCl treatment + 10 mins condition medium and 5 min KCl treatment +20 mins condition medium as indicated in Fig. 1.

Neurons were fixed in 4% PFA at room temperature for 10 minutes after sampling and permeabilized by 70% (v/v) EtOH at 4 degrees overnight. The mouse *Npas4* mRNAs were hybridized with the Quasar^®^ 570 Stellaris RNA FISH Probe set following the manufacturer’s instructions available online. Custom Stellaris^®^ FISH Probes were designed against mouse *Npas4* mRNA by utilizing the Stellaris^®^ RNA FISH Probe Designer (Biosearch Technologies, Inc., Petaluma, CA) available online. We hybridized probes to samples in hybridization buffer (10% Formamide, 10% 20x SSC, 10% Dextran sulfate, 1 mg/mL *Escherichia coli* tRNA, 2 mM Vanadyl ribonucleoside complex and 20 ug/mL BSA) at 37 degree for 4 hours followed by Hoechst staining. Z-stack images were captured on wide-field microscope (DMI4000, Leica) equipped with a CCD camera (DFC365 FX, Leica) and controlled by MetaMorph (Molecular Devices). Objective with NA 1.4 and 63X magnification yielded pixel-size of 146 nm. 35-45 Z-slices were recorded with a 200 nm step-size and 1 second exposure time.

We used FISH-quant [15] to identify and count absolute mRNA numbers and active transcription sites in single cells (Fig. 1). The active transcription sites are detected because nascent mRNAs are transiently attached to the elongating RNA Polymerase II in the gene, accumulating fluorescent probes around active sites, and then appear as highly intense dots (1 or 2, as there are two copies of the gene) in the nucleus of the diploid cell. We and others have confirmed that these nuclear spots mark the active transcription sites because they colocalize in two-color smFISH with probes specific for the gene introns, which are present only in nascent RNAs (data not shown and [18]).

### Monte Carlo sampling and burn-in

The number of iterations (*T*), covariance matrix Σ, and “burn-in” period were determined by monitoring the acceptance rate of proposals and the distribution of parameters and likelihood in the stationary phase of the Monte Carlo algorithm. The rate at which the Markov chain approaches stationarity (i.e. the region with higher likelihood) depends on the covariance matrix Σ used to draw new proposals. We defined the burn-in as the initial period where the log-likelihood was increasing and less than 99.5% of the maximum. The burn-in period is sensitive to the initial parameters and the parameter-stimulus model. Given our experimental data, we verified that *T* = 10^5^ iterations and our covariance matrix Σ were sufficient for BayFish to achieve stationarity and adequately sample the Bayesian posterior distribution after discarding the burn-in. The final covariance matrix Σ was diagonal with 10^*−*5^ for *k*_0_, *k*_1_, *µ*_0_ and 10^*−*3^ for *µ*_1_ proposals. We ran three BayFish replicas for each parameter-stimulus model with a random initial parameter set *θ*.

### Information Criterion and Model Fitting

We used several information criteria, such as the Bayesian Information Criterion [24], Akaike Information Criterion [25], and Deviance Information Criterion [26], to evaluate the likelihood of different models and to penalize model over-fitting.

- *Bayesian Information Criterion*:

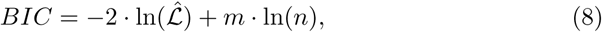
- *Akaike Information Criterion*:

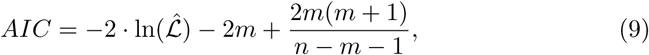

where the maximum likelihood 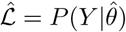 is the maximum value of 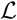 obtained during the BayFish run, *m* is the number of free parameters that were fit, and *n* is the total sample size.

These metrics do not take full advantage of the Bayesian posterior probability estimated by BayFish. To this end, we also used:

- *Deviance Information Criterion*:

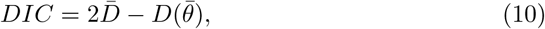

where the deviance is

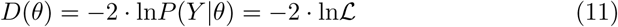

and 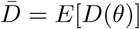 is the mean of the deviance *D*(*θ*) calculated from the Bayesian posterior probability, whereas 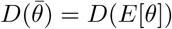 is the deviance of the mean of *θ* calculated from the Bayesian posterior probability.

## Supporting Information

**S1 Fig.**
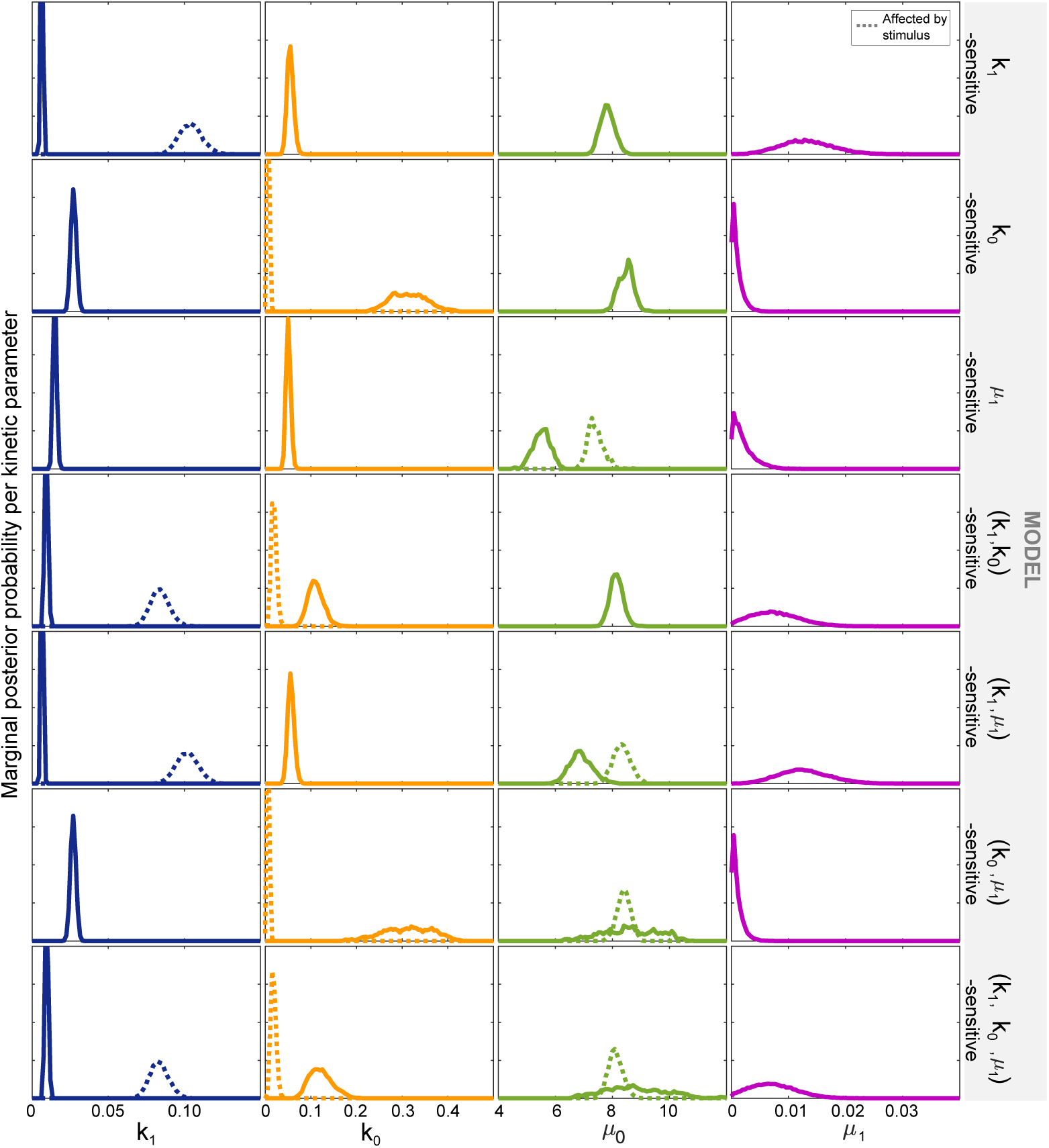
Comparing parameter distributions of different 2-states models. The marginal posterior distribution for each biophysical parameter. Each row corresponds to a different model with the parameter(s) sensitive to stimulus shown in the left. When the parameter is sensitive to stimulus, *uninduced* conditions are shown as continuous lines, *after stimulus* conditions as dotted lines. The results from three MRW replicas were used in all cases.

**S1 Code** can be found at https://github.com/mgschiavon/BayFish.

## Acknowledgments

We are grateful to Sayan Mukherjee and Stefano Di Talia for advice and feedback. This work was supported by a CONACYT graduate fellowship (MGS), the National Institutes of Health Director’s New Innovator Award DP2 OD008654-01 (NEB), the Burroughs Wellcome Fund CASI Award BWF 1005769.01 (NEB), the National Institutes of Health Exploratory/Developmental Research Grant Award R21DA041878 (AEW), and seed funding from the Duke Center for Genomic & Computational Biology (AEW and NEB).

